# Environmental drivers behind the genetic differentiation in mountain chickadees (*Poecile gambeli)*

**DOI:** 10.1101/2023.02.25.529994

**Authors:** P Srikanthan, TM Burg

## Abstract

Anthropogenic climate change has a large impact on wildlife populations and the scale of the impacts have been increasing. In this study, we utilised ddRAD sequence data to investigate genetic divergence and identify the environmental drivers of genetic differentiation between 12 populations of mountain chickadees, family Paridae, sampled across North America. To delineate populations and identify potential zones of hybridisation, we conducted a discriminant analysis of principal components (DAPC), admixture analysis, and calculated pairwise Fst values. The DAPC revealed four clusters: southern California, eastern Rocky Mountains, northwestern Rocky Mountains and Oregon/northern California. We then used BayeScEnv to highlight significant outlier SNPs associated with the five environmental variables. We identified over 150 genes linked to outlier SNPs associated with more than 15 pathways, including stress response and circadian rhythm. We also found a strong signal of isolation by distance. Local temperature was highly correlated with genetic distance. Maxent simulations showed a northward range shift over the next 50 years and a decrease in suitable habitat, highlighting the need for immediate conservation action.

## Introduction

Anthropogenic climate change is increasingly affecting ecosystems worldwide. The steady increase in temperature means and variability can affect the functioning of organisms and alter ecosystems that have existed on this planet for millennia. While most ecosystems are sensitive to changes in the environment, mountain ecosystems are particularly vulnerable to environmental changes (Beniston, 2003). Owing to increasing temperatures and reduced snow cover, organisms in mountainous regions may shift their elevational range to higher altitudes, potentially leading to population fragmentation and extinction (Calkins et al., 2012; McDonald & Brown, 1992; Parmesan, 2006; Wilson et al., 2007). Climate change also significantly affects phenology, such as the timing of flowering or breeding (Walther et al., 2002). The extent and effects of these changes are a current area of study, and the pace at which changes occur requires commensurate efforts in the form of conservation research to prevent the breakdown of ecosystems and extinction of organisms (Christmann & Menor, 2021; Payne et al., 2020).

The North American landscape is full of physical barriers that affect species dispersal, and therefore causing genetic divergence (Antonelli, 2017; Boutilier et al., 2014; Machado et al., 2018). In addition, the wide range of environmental conditions across the continent makes it an ideal area to study the effect of the environment on genetic differentiation. In particular, species with low migration rates could experience significant divergence (Keyghobadi et al., 1999). Therefore, it is crucial to study the population genetics of these species to investigate their resilience to climate change (Hanski et al., 2006). Despite being able to traverse long distances, birds are unable to pass through certain barriers, makes them a good model for studying the effects of physical barriers on divergence (Greenwood & Harvey, 1982).

The mountain chickadee, *Poecile gambeli*, is found across western North America and primarily occupies coniferous forests. As many as seven subspecies have been described and are divided into two mtDNA groups, eastern and western, with limited contemporary gene flow observed between them (Hindley et al., 2018; Spellman et al., 2007). A recent study using microsatellite data identified Washington as a potential contact zone between two clades (Hindley et al., 2018). Ubiquitous across the west, the mountain chickadee has limited migratory capabilities, and high philopatry makes it an ideal candidate for studying the effects of geography and climate on genetic differentiation.

In this study, we examined the population structure of mountain chickadees from populations across the range using double digest restriction-site associated DNA, ddRAD, data. In addition, we aimed to answer the following questions: (1) What are the environmental drivers of genetic differentiation? Previous studies using mtDNA have shown that isolation by distance influences differentiation (Hindley et al., 2018); however, more information is needed on the effects of other variables such as temperature and precipitation. (2) Which genes or pathways are undergoing selection across populations? Given the pace of climate change, it is essential to investigate the pathways undergoing selection to identify gene-environment interactions. (3) How much will the changing climate affect the habitat of mountain chickadees over the next 50 years? Although we expect a northward shift in habitat or a shift in elevational range, understanding how a common species is affected by climate change can provide valuable insights to inform conservation decisions on other co-distributed species.

## Methods

### Sampling and DNA extraction

A total of 94 mountain chickadee samples were collected from 12 locations (Figure 1, Table 1). Birds were caught using mist nets and either blood or feather samples were collected. Following Aljanabi et al. (1997), we used ~40 ul of blood to extract DNA from samples. Library preparation for ddRAD sequencing was done at the University of Laval using the enzymes *Pstl*, *Nsil*, and *Mspl* and sequencing was done at Genome Quebec on an Illumina NovaSeq 6000 S4 PE100.

**Figure 1.**
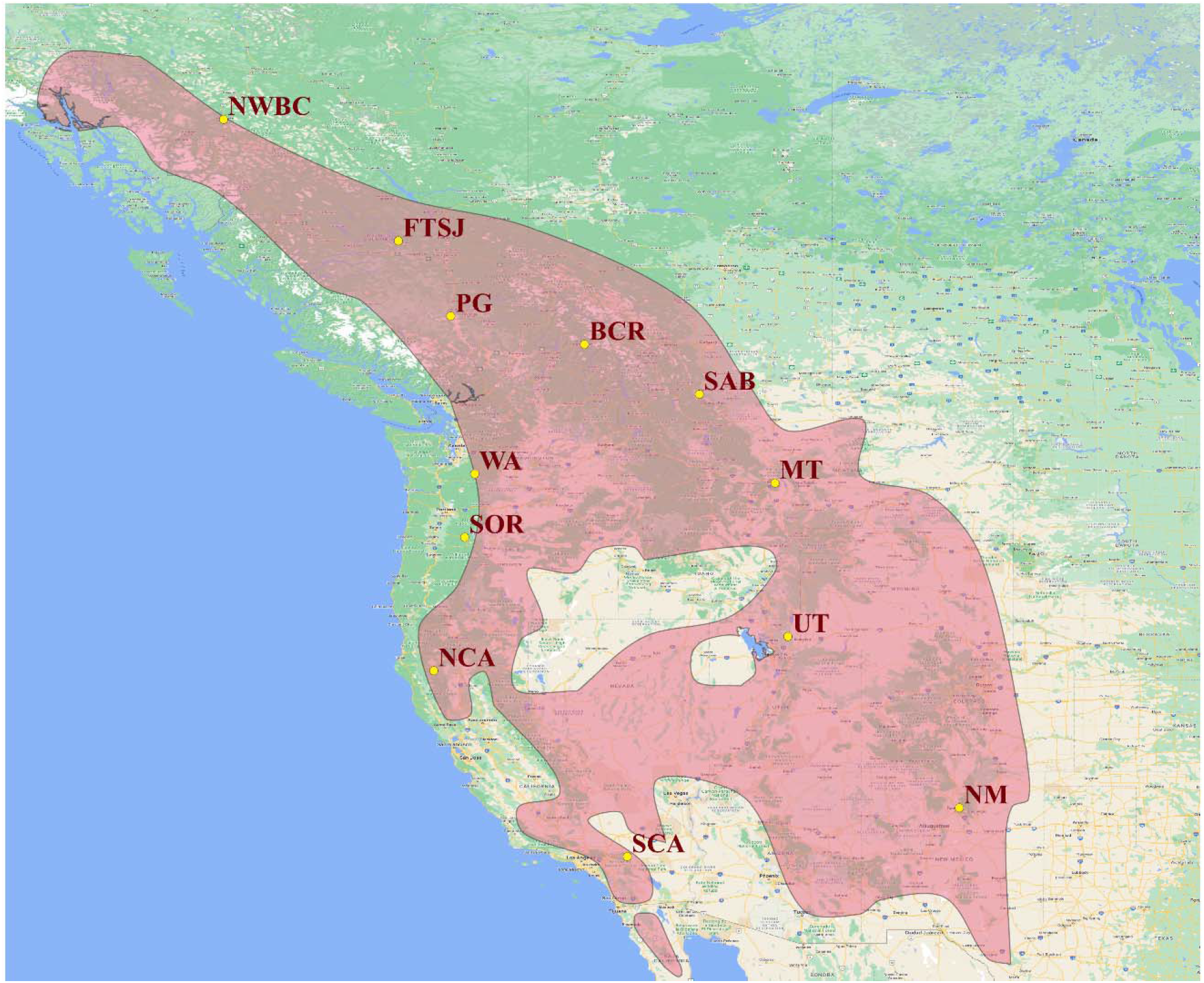
Mountain chickadee sampling sites in western North America along with the range map from the IUCN Red List (pink)

**Table 1.** List of sampling sites and sample sizes for mountain chickadees used in this study. See Fig 1 for location.

| Location | Abbreviation | Sample size |
| --- | --- | --- |
| Revelstoke, BC | BCR | 4 |
| Fort St James, BC | FTSJ | 8 |
| Northwest British Columbia | NWBC | 2 |
| Prince George, BC | PG | 2 |
| Washington | WA | 8 |
| Montana | MT | 7 |
| Utah | UT | 10 |
| Southern Alberta | SAB | 2 |
| Southern Oregon | SOR | 10 |
| Northern California | NCA | 5 |
| Southern California | SCA | 3 |

### Bioinformatic pipeline

Sabre was used to demultiplex the ddRAD data. Adapters were removed and fastq files were trimmed to 80 bp using Cutadapt (Martin, 2011). The fastq files with the trimmed sequences were aligned to a black-capped chickadee reference genome provided by Scott Taylor (University of Colorado, Boulder) with BWA-MEM using default settings (Branch et al., 2022; Li & Durbin, 2009; Wagner et al., 2020). The resulting bam files were used to identify SNPs with gstacks. The gstacks output was used to create a vcf file and calculate summary statistics with populations using the default parameters while limiting the SNPs to one per locus (Catchen et al., 2013). SNPs were subsequently filtered using Vcftools to keep bi-allelic SNPs with a minor allele frequency ≥ 0.05 (Danecek et al., 2011). SNPs with > 50% missing data and individuals with > 30% missing data were excluded. The filtered vcf files contained 28,600 sites and 61 individuals from 11 populations; no individuals from NM remained after filtering.

### Population analyses

To analyse the population structure, we conducted a DAPC (discriminant analysis of principal components) in R using the packages ‘adegenet’,’ape’, ‘vcfR’, and ‘ade4’ (Dray & Dufour, 2007; Jombart, 2008; Knaus & Grünwald, 2016; Paradis & Schliep, 2018). DAPC accounts for both within- and between-group differences when clustering individuals. After clustering, we plotted the linear discriminants in 2d and 3d plots using the packages ‘ggplot2’ and ‘plotly’ (Sievert, 2020; Wilkinson, 2011).

To identify the number of ancestral populations, we used ADMIXTURE with default settings iteratively for K=1-7 (Alexander et al., 2009). Linkage disequilibrium pruning and vcf-to-bed conversion were performed using PLINK (Purcell et al., 2007).

Pairwise Fst values between populations were calculated with Arlequin using the default settings (Excoffier & Lischer, 2010). Vcf-to-arp conversion was done using PGDSpider2 (Lischer & Excoffier, 2011).

### Outlier loci and environmental variation

To identify the candidate loci under selection, we used BayeScan (Foll & Gaggiotti, 2008). BayeScan is a Bayesian algorithm based on the multinomial-Dirchlet model, which identifies loci under selection using differences in allele frequencies between populations. We set the pr_odds value at 350, with a total of 1,00,000 iterations. The pr_odds value defines the likelihood of the neutral model compared with the selection model.

To identify divergence associated with environmental variation, we used BayeScEnv for five environmental variables: temperature, precipitation, altitude, temperature seasonality and precipitation seasonality (Villemereuil & Gaggiotti, 2015). BayeScEnv, which is based on the F-model, detects local adaptation linked to a given environmental variable using Bayesian methods. In this study, we set the pr_jump value at 0.05 with a total of 50,000 runs. The pr_jump value is similar to the pr_odds value used in BayeScan. Using the ‘sp’ and ‘raster’ packages, we extracted 30s resolution from https://www.worldclim.org/ for the following environmental variables: Annual Mean Temperature (BIO1), Temperature Seasonality (stdev×100)(BIO4), Annual Precipitation (BIO11), Precipitation Seasonality (Coefficient of Variation) (BIO11), and Elevation from the SRTM data (Bivand et al., 2013; Fick & Hijmans, 2017; Hijmans, 2020). The environmental data were standardised before the BayeScEnv analysis.

To test for isolation by distance and the effect of the above environmental variables on genetic differentiation, we conducted Mantel and partial Mantel tests using GenAlEx and the R package ‘vegan’ (Dixon, 2003; Mantel, 1967; Peakall & Smouse, 2012). Mantel tests are commonly used in population genetics to test for correlation between two matrices, while partial Mantel tests use a third matrix to account for another variable (Diniz-Filho et al., 2013).

### Genes of interest

We used the gff file for black-capped chickadees provided by Scott Taylor to identify genes under selection and investigate gene–environment interactions. We ran a custom R script to identify genes present within 100 kb of the outlier loci from the BayeScan/BayeScEnv analyses. Subsequently, to identify the pathways and functions associated with these genes, we ran a gene ontology with ShinyGO using a human and zebra finch gene set. The genes were also analysed individually (Ge et al., 2019).

### Species distribution models

To understand the distribution of the species in the past and future, we created species distribution models for the current distribution, the last glacial maxima, and four Representative Concentration Pathways (RCP) in 2050 and 2070. RCPs describe different climatic futures depending on the emission of greenhouse gases with the four main RCPs being RCP2.6, RCP4.5, RCP6, and RCP8.6 (van Vuuren et al., 2011).

We used the Maxent algorithm in the ‘Wallace’ package (Kass et al., 2018). This algorithm uses occurrence data in conjunction with environmental layers to provide a potential distribution for a species. Presence data were obtained from iNaturalist observations over a 13 year period from 2010-2022 (GBIF.org, 2022) which we spatially thinned using a 10 km threshold. These points were then partitioned into six clusters using the random k-fold method (Aiello-Lammens et al., 2015). Using these 5330 points, we created a SDM with 14 environmental variables.

We tested 12 models: linear (L), quadratic (Q), hinge (H), and linear-quadratic-hinge (LQH), with a regularization multiplier between 1-3 using a step value of 1. Model selection was done using AICc and AUC values (Kass et al., 2021; Phillips, 2017). The trained models were projected onto the LGM and RCP scenarios using environmental layers obtained from http://www.paleoclim.org and CCSM4, respectively using a 10% minimum training presence threshold (Brown et al., 2018; Hill, 2015; Karger et al., 2017; Pisias & Moore, 1981).

## Results

### Population structure

We looked at over 28,500 loci in 61 individuals from 11 populations after filtering. Prior to filtering, there were over 1,000,000 SNPs for 94 individuals from 12 populations. The DAPC, consistent with Hindley et al. (2018), revealed four distinct clusters: southern CA; northern CA/southern OR; eastern Rockies (MT, UT, SAB); and northwestern Rockies (WA, BCR, FTSJ, PG, NWBC) (Figure 2a). The first three linear discriminants (LD) explained 98.5% of the variance, while the fourth explained approximately 1.5%. Pairwise LD plots (Figure 2b) accounted for the effect of altitude on genetic differentiation.

**Figure 2.**
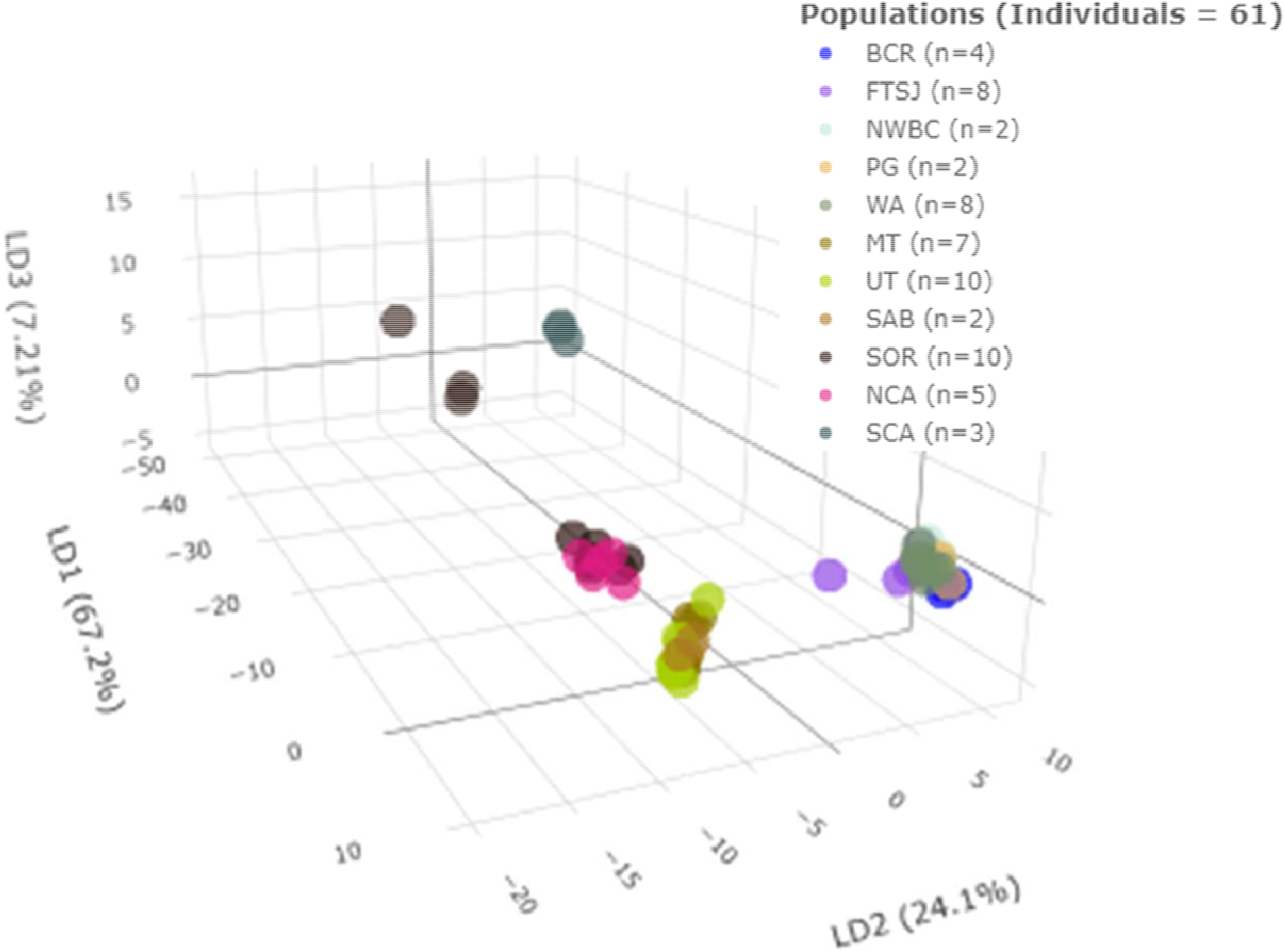
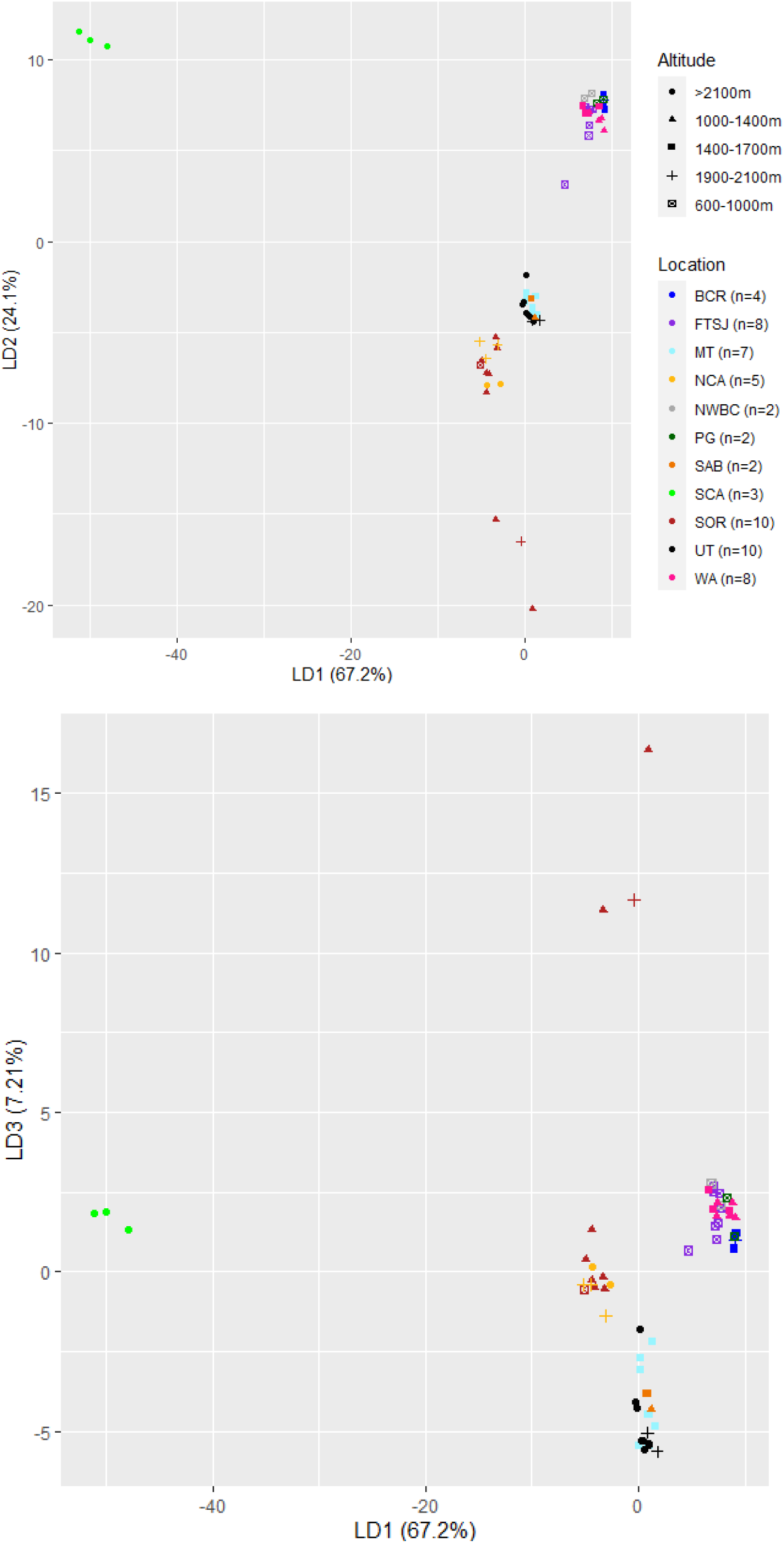
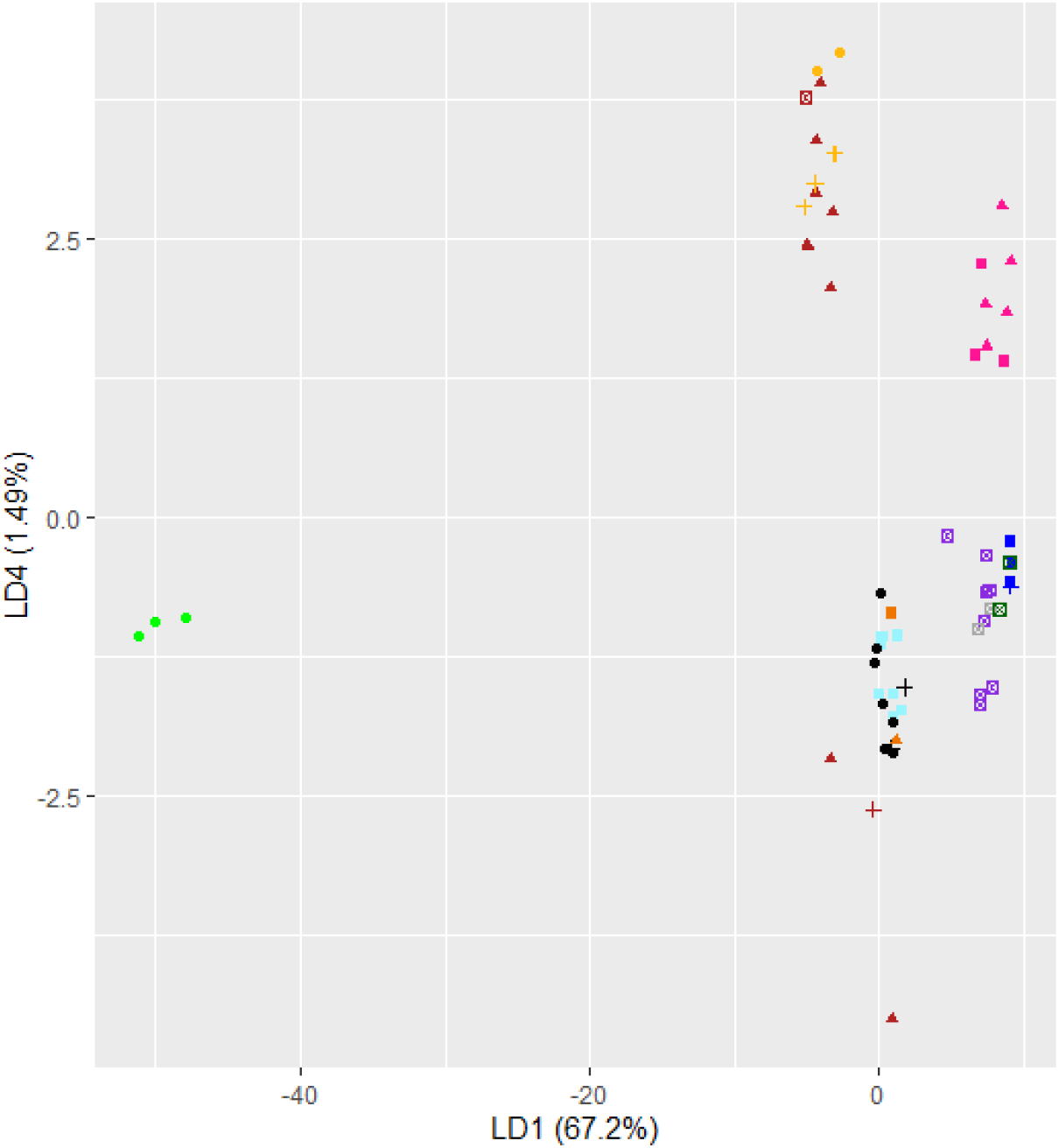
(a) 3d DAPC plot showing four distinct clusters(b) LD1 vs LD2 (c) LD1 vs LD3 (d) LD1 vs LD4

As ADMIXTURE does not predict the K value, we ran the analysis for K=1-7 (Figure 3a-b). Figure 3(b) shows the CV errors for each K value, where K=1-3 are strongly supported (CV error 0.668-0.696). K=4-6 coincide with the DAPC despite having a slightly higher CV error (0.758-1.026). For K=4, the SCA population splits from the SOR/NCA population to form a separate cluster consistent with the DAPC analysis. At K=5 WA separates from the other northwestern Rocky populations and BCR individuals and three FTSJ individuals show admixture with WA.

**Figure 3.**
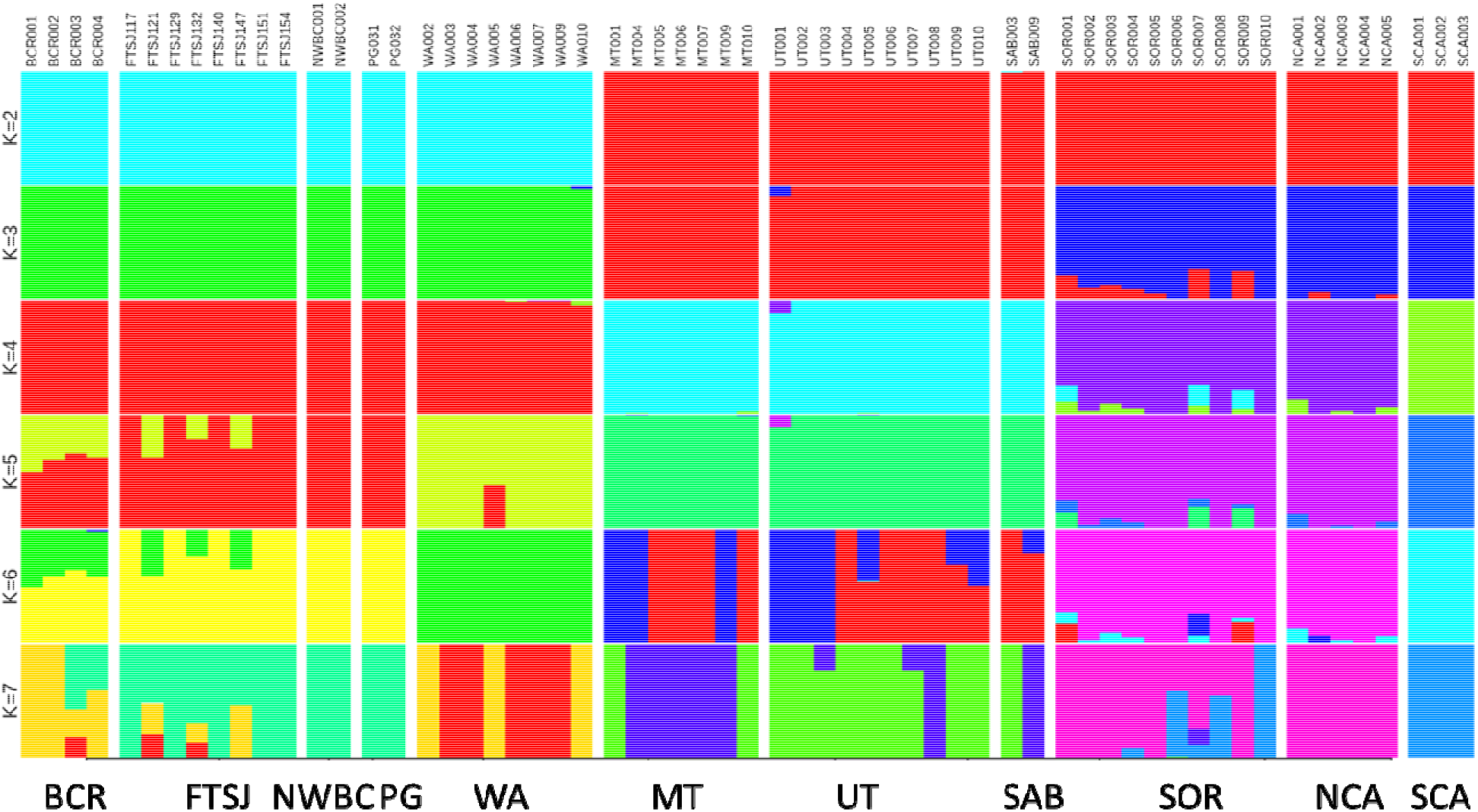
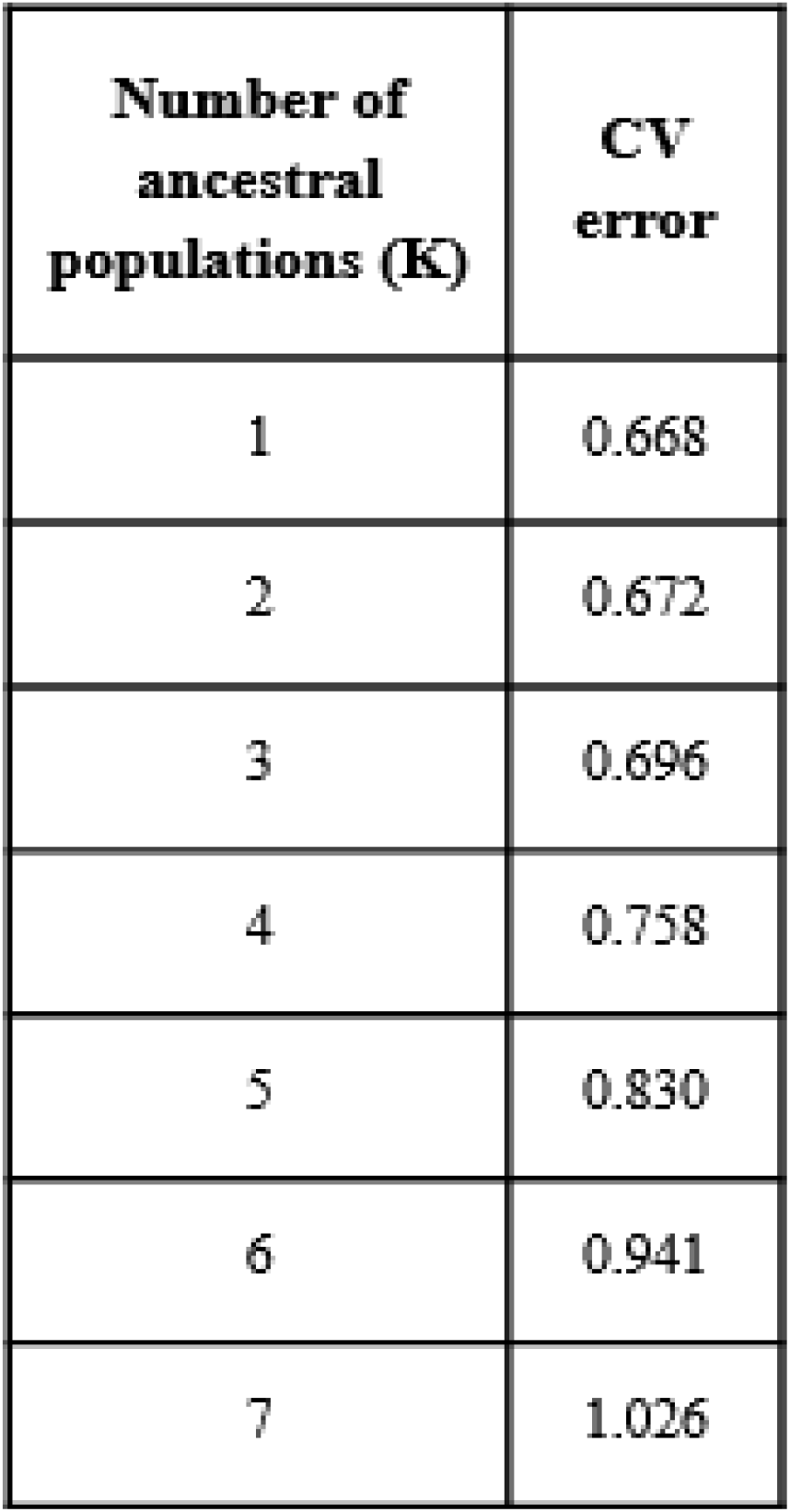
(a) Admixture plot for K=2-7 (b) Cross validation errors for K=1-7

The pairwise Fst values, consistent with the DAPC, showed four broad clusters, with the SCA population having the highest Fst values between every pair (Figure 4). The p-values were significant for most cases, except those that included NWBC, PG, or SAB populations. Within the northwestern Rockies group, WA was significantly different from all of the other populations except PG. Pairwise Fst values ranged from 0.02 to 0.37, the highest significant value is seen between BCR-SCA while the smallest are between the BCR-FTSJ and UT-MT populations.

**Figure 4.**
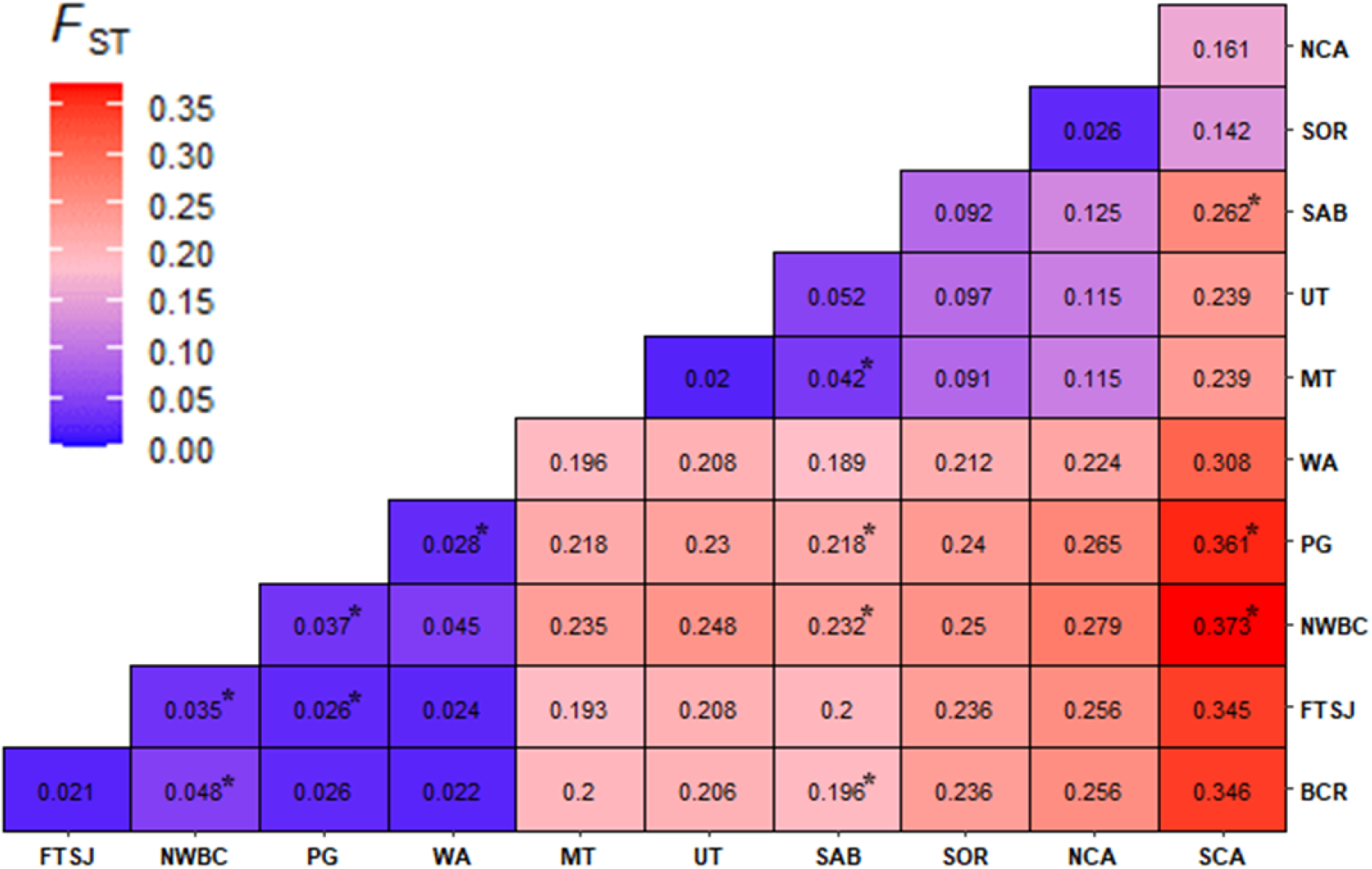
Pairwise Fst values for 11 populations. Asterisks indicate non-significant observations (p-values >0.05).

### Outlier loci and gene-environment analysis

To identify areas of genomic divergence and their association with environmental factors, we used BayeScan and BayeScEnv, two commonly used genome scan programs (Foll & Gaggiotti, 2008; Villemereuil & Gaggiotti, 2015). Outlier loci associated with temperature (Temp), altitude (Alt), precipitation seasonality (PS), temperature seasonality (TS), and precipitation (Prec) were identified using BayeScEnv, whereas BayeScan was used to identify all possible outliers, regardless of their association with environmental variables.

We identified 2251 outlier loci at a false discovery rate (FDR) of 0.0001 using BayeScan, despite using a conservative model with a pr_odds value of 350 (Figure 5a). Several SNPs had a false discovery rate of 0, which was converted to 10e^-10^ for visualisation. Similarly, the BayeScEnv analysis, with an FDR of 0.0001, showed 1564 outlier SNPs associated with temperature, 2060 with temperature seasonality, 805 with precipitation, 1090 with precipitation seasonality, and 1606 with altitude (Figure 5b-f). Several SNPs had a false discovery rate of 0, which was converted to 10e^-6^ for visualisation. We used different values for visualisation between the BayeScan and BayeScEnv analysis due to several SNPs having FDRs of 10e^-6^ in the former.

**Figure 5.**
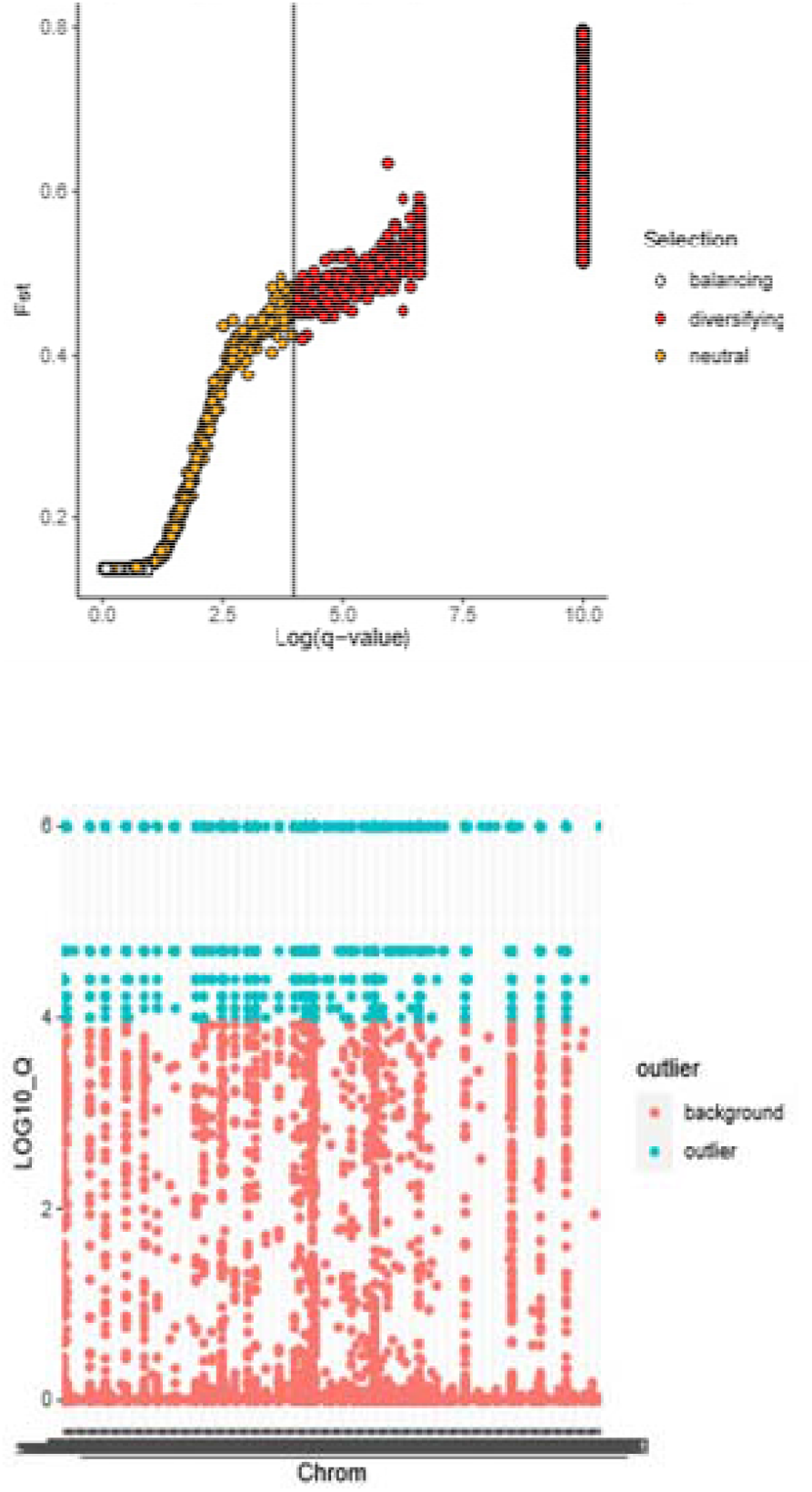
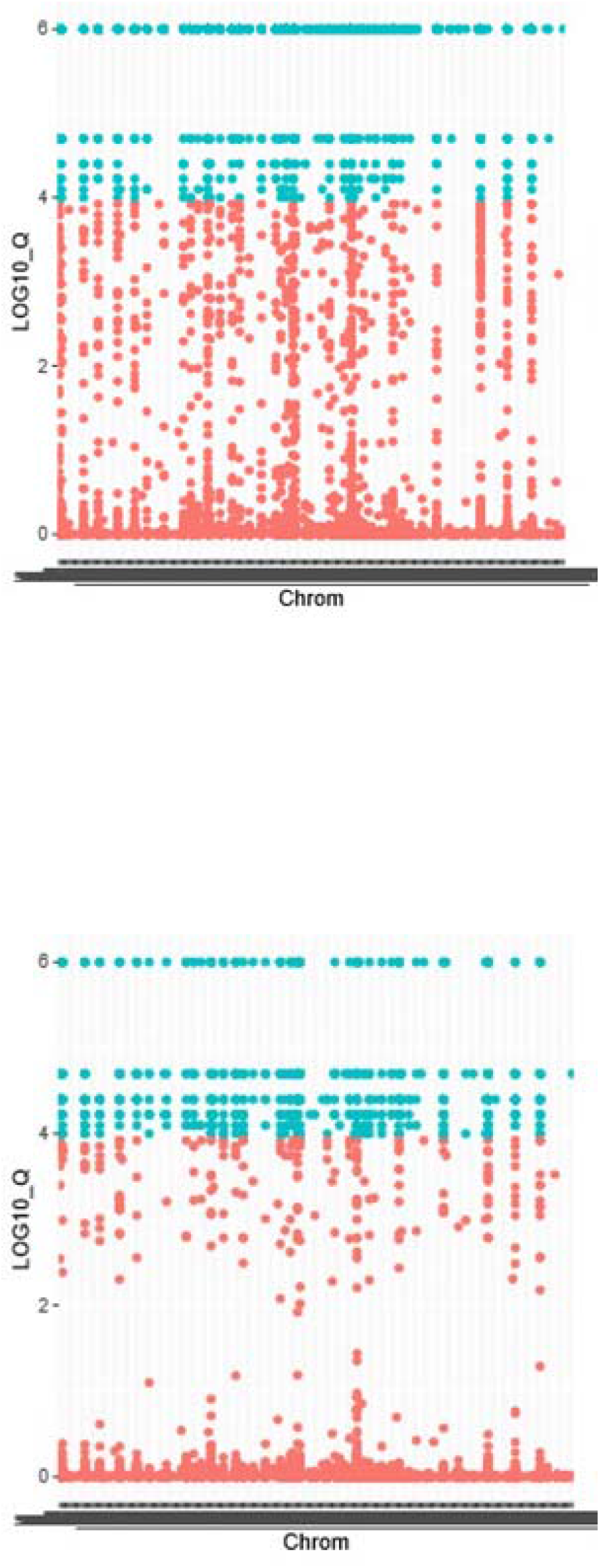
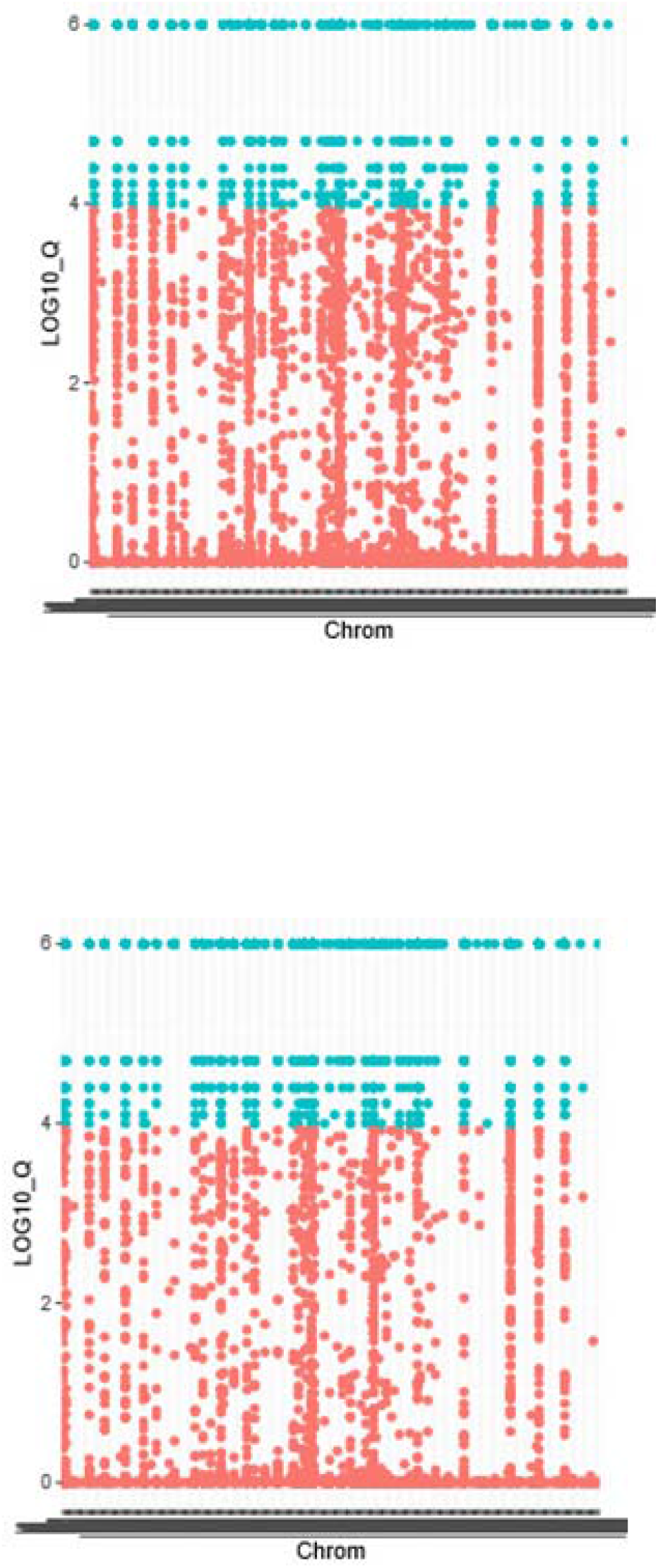
BayeScan and BayScEnv plots with correlation q-values for genetic divergence where q-value = −log(FDR). (a) BayeScan plot (N=2251, FDR=0.001). (b) BayScEnv Temperature plot (N=1564, FDR=10^-4^). (c) BayScEnv Temperature Seasonality plot (N=2060, FDR=10^-4^). (d) BayScEnv Precipitation plot (N=805, FDR=10^-4^). (e) BayScEnv Precipitation Seasonality plot (N=1090, FDR=10e^-4^). (f) BayScEnv altitude plot (N=1606, FDR=10e^-4^).

To test whether the large number of outlier loci was due to the isolated SCA population, we ran the same tests for pairs of the identified DAPC clusters and with all populations except for SCA. We found a significant decrease in the number of outlier loci identified in the pairwise cluster tests. The decrease differed amongst clusters, but the number of outlier loci ranged from 4-800. In the case of precipitation, the outliers ranged from 30-500 compared to 805. However, there was no change when we repeated the analysis with every population excluding the SCA population.

We performed Mantel and partial Mantel tests to investigate the possibility of isolation by distance and the effect of the aforementioned environmental variables on divergence. Geographic distance (R_xy_= 0.54) was most highly correlated with genetic distance, followed by temperature (R_xy_= 0.40) (Figure 6a-b). All the results were statistically significant (p < 0.0003). The partial Mantel tests highlighted a similar pattern; however, when the z-axis was either temperature, temperature seasonality, or geographic distance; the results for precipitation seasonality did not meet the cut-off criterion of p = 0.05. In addition, the results of temperature seasonality when the z-axis was geographic distance did not meet the cut-off criteria.

**Figure 6.**
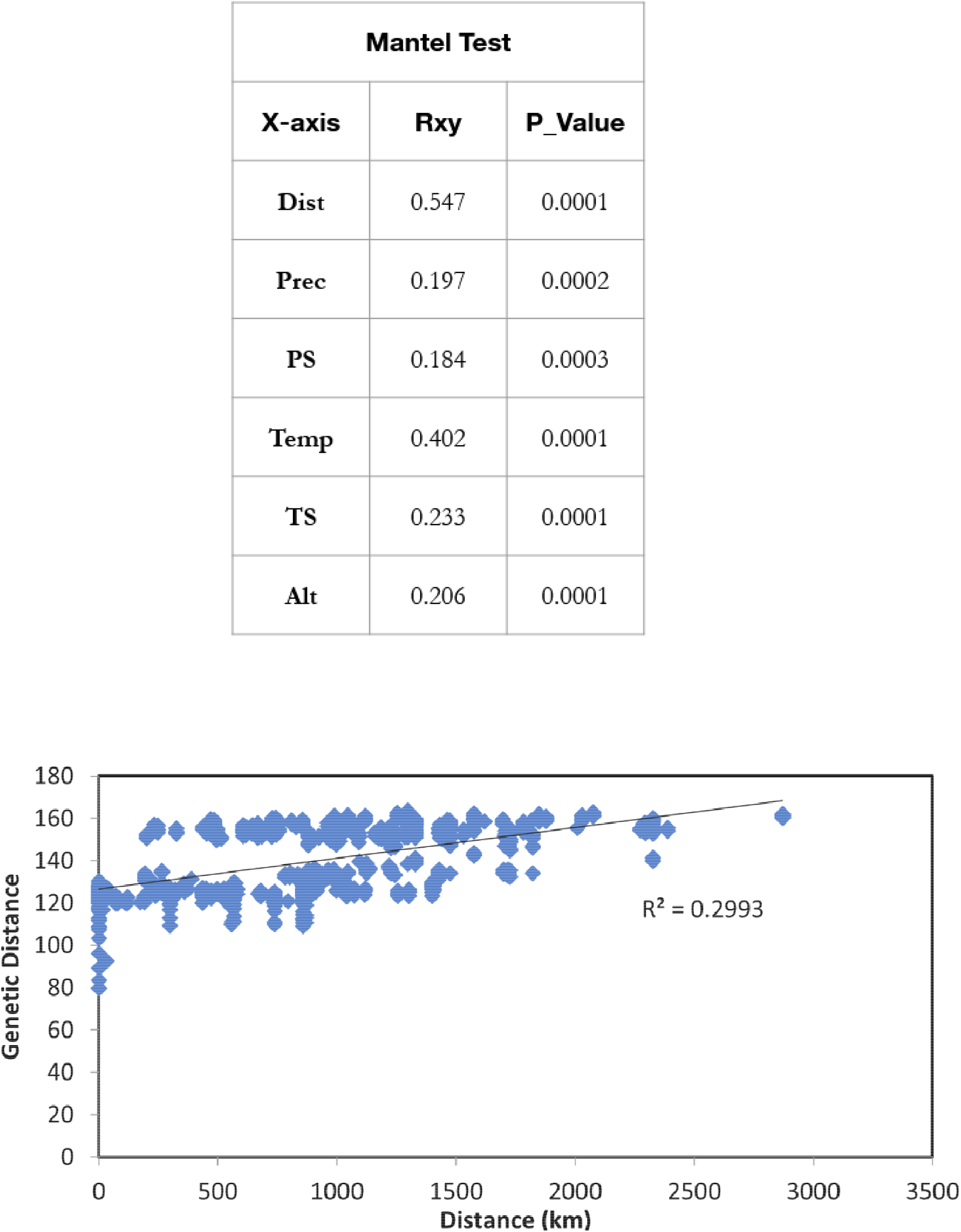
(a) Mantel test R_xy_ and p-values. Geographic distance-Dist, Precipitation-Prec, Precipitation seasonality-PS, Temp-Temperature, Temperature Seasonality-TS, Altitude-Alt (b) Geographic vs genetic distance plot.

### Genes of interest

We identified 181 genes within 100 kb of temperature-associated SNPs and 189 genes from the BayeScan outlier SNPs. Genes of interest were analysed only for these two cases because of the overlap in SNPs across environmental variables. ShinyGO identified 15 significant pathways (Supplementary, gene_SuppInfo.xlsx) for the BayeScan genes, which included thermogenesis (p = 4.58e^-5^).

Subsequently, we grouped the genes based on their biological functions using ShinyGO (Ge et al., 2019). The BayeScan analysis revealed 37 genes associated with response to stress, 22 genes related to immune system processes, and three genes associated with circadian rhythm. CSNK1D, a clock gene, was one of the genes associated with circadian rhythm, which has been shown to play a role in migration (Steinmeyer et al., 2009). In addition, several genes affecting immune response, response to external stimuli, and growth were found to be associated with outlier SNPs. We also identified genes associated with the reproductive system in all the analyses. The JAM3 gene is known to play a role in spermatogenesis and the RNF212 gene has been shown to influence meiotic recombination. A similar grouping of genes was observed using the BayeScEnv temperature-associated SNPs, with the number of genes associated with each function differing.

We obtained similar results when using the zebra finch genome, although the number of unmapped genes was higher than that in the human genome (BayeScan – 137 vs 87 unmapped, BayeScEnv – 108 vs 64 unmapped).

### Species distribution models

We created species distribution models to understand the effects of climate change on mountain chickadee habitat (Figure 7). We ran 12 models with a regularisation multiplier between 1-3 with a step value of one. While all models had an AUC value of 0.93 or above, the hinge model with regularisation multiplier 1 was chosen as the best model based on AICc and AUC values.

**Figure 7.**
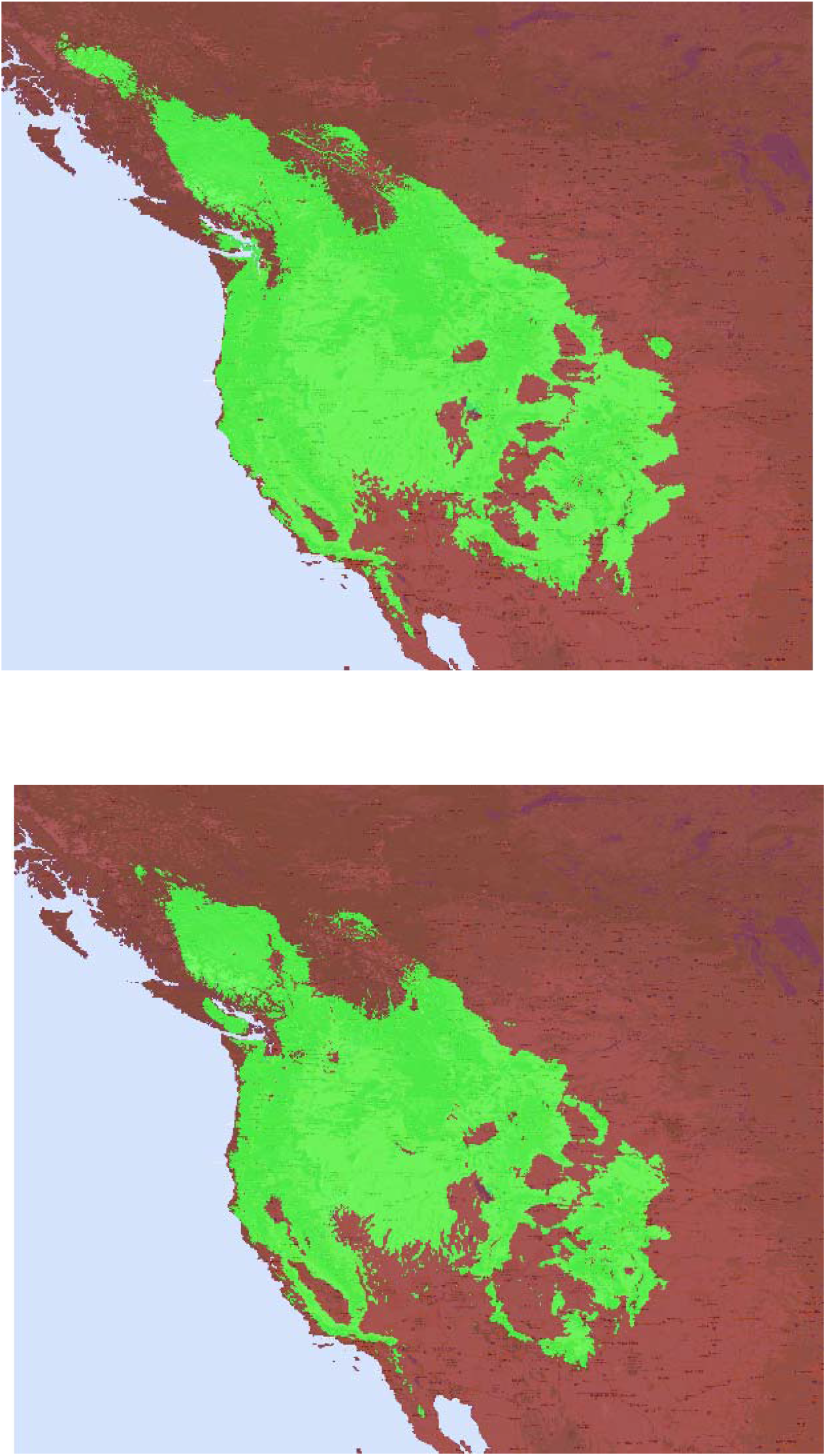
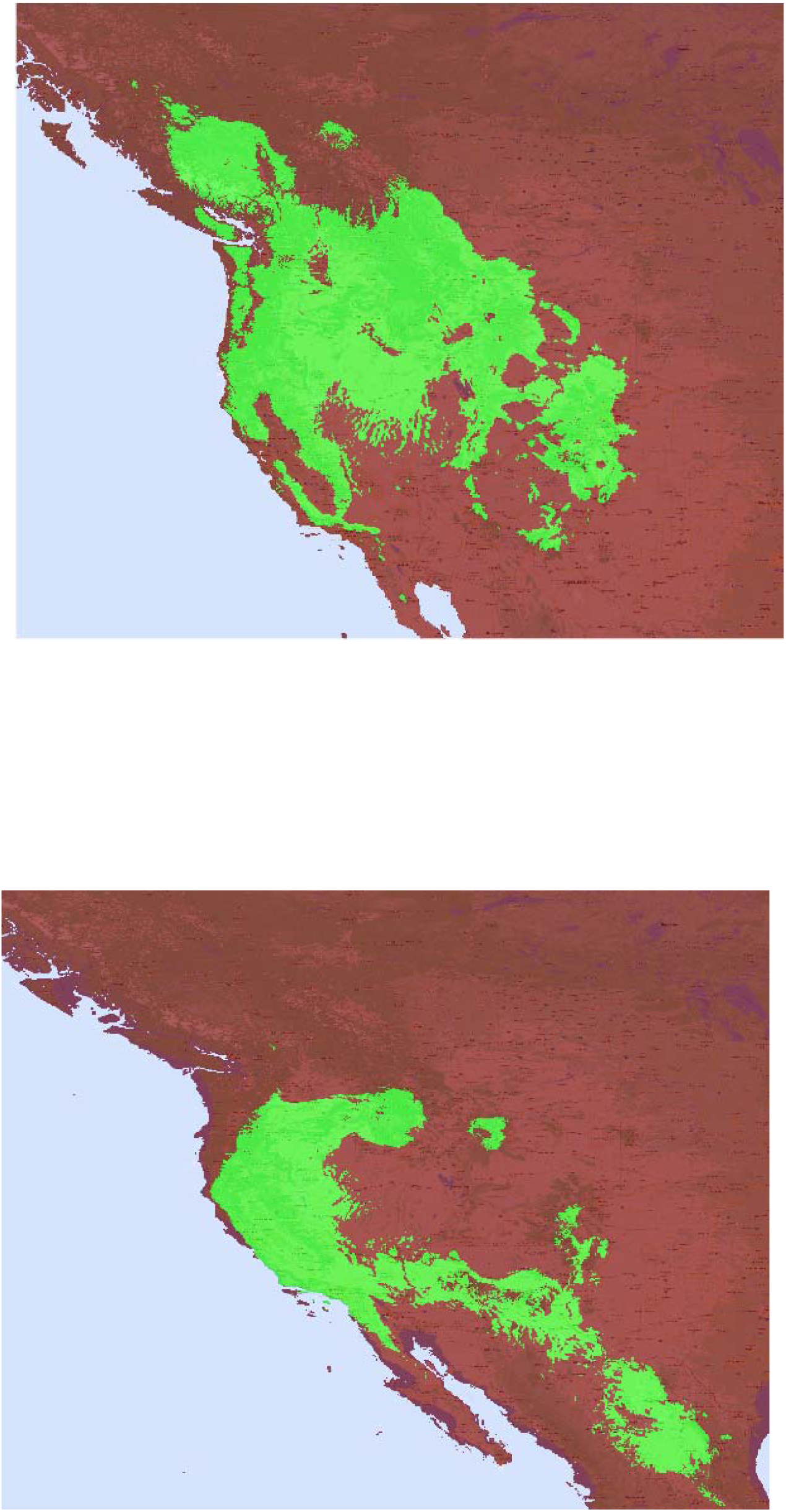
Species distribution models of mountain chickadees using the 10^th^ percentile training presence threshold. Legend: green-presence, brown-absence (a) Current distribution. (b) 2050 distribution under RCP 8.5 (c) 2070 distribution under RCP 8.5. (d) Distribution during the last glacial maxima 21 kya.

The SDMs show a significant decrease in suitable habitat over the next five decades across all RCP scenarios except RCP2.6, which is an ideal projection of future climate. In addition, a northward shift in habitat is also observed. The SDM for the last glacial maximum is consistent with previous studies, with populations present near the coast and extending into Mexico (Manthey et al., 2012) (Figure 7d). The SDMs for other RCPs, response curves, and model statistics are available in the supplementary material.

## Discussion

In this study, apart from delineating population structure, we aimed to answer the following questions: (1) What are the environmental drivers of genetic differentiation? (2) Which genes or pathways are undergoing selection across populations? (3) How much will climate change affect the habitat of mountain chickadees over the next 50 years?

DAPC revealed four primary clusters, which coincided with our ADMIXTURE and pairwise Fst values. The eastern Rocky Mountain cluster (UT, MT, SAB) is separated from the northwestern Rocky Mountain populations by the Rocky Mountains. These populations could have diverged because of the presence of physical barriers or differences in habitat on either side of the mountains. This is evident from the high, but not significant, pairwise Fst values between the BCR and SAB populations, which are geographically close (300 km), but separated by the Rockies. In addition, the SCA population is of particular interest because it is genetically isolated from all other populations. As indicated by the Mantel tests and previous studies, this could be due to its distance from other populations or due to geographic/environmental features within its habitat (Spellman et al., 2007; Hindley et al., 2018).

Our outlier SNP analysis revealed several loci under selection associated with environmental conditions. Despite the use of conservative models, we identified several outlier loci highlighting the genetic diversity of the species. However, because the large number of loci could have been due to the isolated SCA population, we conducted the same analyses by (1) excluding the SCA population and (2) between the DAPC clusters. There were no significant differences in the first case, whereas we observed a decrease in the second case. This decrease can be attributed to the low genetic distances between populations within the same cluster. Additionally, the overlap of SNPs across analyses and the existence of several SNPs with an FDR of zero highlights the genetic diversity present in the species. Divergence across populations is expected because of the reduced gene flow among them due to the non-migratory nature of the species (Templeton, 2006; Eckert et al., 2008; McCallum et al., 2020).

Consistent with mtDNA and microsatellite studies, genetic diversity is highly influenced by geographic distance (Hindley et al., 2018). The strong correlation between geographic and genetic distance (R_xy_ = 0.55) indicates isolation by distance. However, we cannot discount the role of habitat differences, given that local temperature (R_xy_ = 0.4), temperature seasonality (R_xy_ = 0.23), and altitude (R_xy_ = 0.2) are also significantly correlated with genetic distance. The rapid increase in global temperatures could affect the genetic isolation in the coming years. Additionally, an increase in temperature forces species to shift their elevational range. Precipitation and precipitation seasonality are predicted to increase with rising temperatures, resulting in more extreme climate scenarios (Boer, 2009; Pendergrass et al., 2017). As a result, despite their weak correlation with divergence, these factors could play a major role in the future of these species.

We identified genes associated with SNPs undergoing selection. Over 30 genes linked to stress response were found to be near with SNPs associated with temperature. We also found over 19 genes linked to response to external stimuli, and ShinyGO analysis revealed that the thermogenesis pathway had a significant number of genes involved. Previous studies have shown that increasing temperatures are linked to the activation of the stress response in birds in the form of thermoregulatory strategies such as panting and increased glucocorticoid levels (Bohler et al., 2021; Mentesana & Hau, 2022; Siegel, 1980). This is of particular concern because of rising global temperatures, which could lead to negative consequences for the species. In addition, the selection of a clock gene, CSNK1D, could imply a change in phenology in response to rising temperatures (Milligan et al., 2009).

We observed a northward shift in suitable habitat over the next five decades with the SDM. This pattern was observed for all RCP scenarios, except for RCP2.6. However, RCP2.6 is an ideal scenario where all expected climate change goals are fulfilled, and temperatures increase by 1°C above pre-industrial levels by the year 2050 and remain the same in 2070 (van Vuuren et al., 2011). While shifting to cooler habitats is a normal thermoregulatory response, the massive decrease in suitable habitats for such a common species is worrying (Siegel, 1980). Additionally, because the model does not account for other factors, such as human-induced habitat loss, competition, and invasion; the amount of suitable habitat could be much less than predicted. This could lead to a further population decline in this species.

## Conclusions

Mountain chickadee genetic distance is highly correlated with geographical distance and temperature. Genes affecting several essential functions associated with outlier SNPs were identified, highlighting the genetic diversity and selection pressure faced by the species. The identification of genes related to circadian rhythm may underlie changes in phenology. In addition, the large decrease in suitable habitat over the next five decades for a common species highlights the need for immediate action to protect this species and other species from extinction.

## Acknowledgements

We thank NSERC Discovery Grant, Alberta Innovates and Mitacs for funding and Burke Museum of Natural History, University of Michigan Museum of Natural History, University of California, Berkeley Museum of Vertebrate Zoology, Louisiana State University Museum of Natural Science, and Smithsonian Museum of Natural Science for providing some of the samples used in this study. Samples were also collected by members of the Burg Lab at the University of Lethbridge including John Hindley, Brendan Graham, Linda Lait, Paulo Pulgarin, Kim Dohms and numerous field assistants. We also thank the National Park Service, USFWS, Parks Canada and Canada Wildlife Service for their assistance.

## Data accessibility and benefit sharing

### Data accessibility

Multiplexed ddRAD sequence data will be uploaded to a data repository prior to publication.

### Benefit sharing

A research collaboration was developed between researchers from India and Canada with funding from Mitacs. Our research also addresses a priority concern, climate change, and provides benefits by the sharing of our data and results on public databases.

## Author contributions

The authors confirm contribution to the paper as follows: Study conception and design: Srikanthan and Burg. Data collection: Burg. Data analyses: Srikanthan. Wrote the paper: Srikanthan and Burg. All authors reviewed the results and approved the final version of the manuscript.

