## Supplemental_Data for "Environmental drivers behind the genetic differentiation in mountain chickadees (*Poecile gambeli)*": Supplemental_Info_README.docx

Srikanthan P, TM Burg

**Table of Contents:**

| **Legend for supplementary files** | Page 2 |
| --- | --- |

**gene_SuppInfo.xlsx**

This supplement contains information on the genes and pathways identified using ShinyGO.

| **Sheet Name** | **Information** |
| --- | --- |
| Temperature_Human | Genes identified using BayeScEnv outlier SNPs associated with temperature, using a human reference on ShinyGO |
| Temperature_Finch | Genes identified using BayeScEnv outlier SNPs associated with temperature, using a Zebra Finch reference on ShinyGO |
| Bayescan_Human | Genes identified using BayeScan outlier SNPs, using a human reference on ShinyGO |
| Bayescan_Finch | Genes identified using BayeScan outlier SNPs, using a zebra finch reference on ShinyGO |
| Bayescan_Pathways_Human | Pathways and genes identified using BayeScan outlier SNPs, using a human reference on ShinyGO |

**Fig7_SuppInfo.zip**

This file contains all data related to the species distribution model.

The ‘Maps’ folder contains all *.TIF files of the range maps for every RCP from RCP2.6 to RCP8.5 for both 2050 and 2070.

The ‘Model_Statistics’ folder contains all files related to model selections and the response curves.

‘Poecile_gambeli_Points.xlsx’ contains the spatially thinned datapoints used for the range maps.
