## Supplementary figures and images for "Environmental drivers behind the genetic differentiation in mountain chickadees (*Poecile gambeli)*"

### auc_diff.png

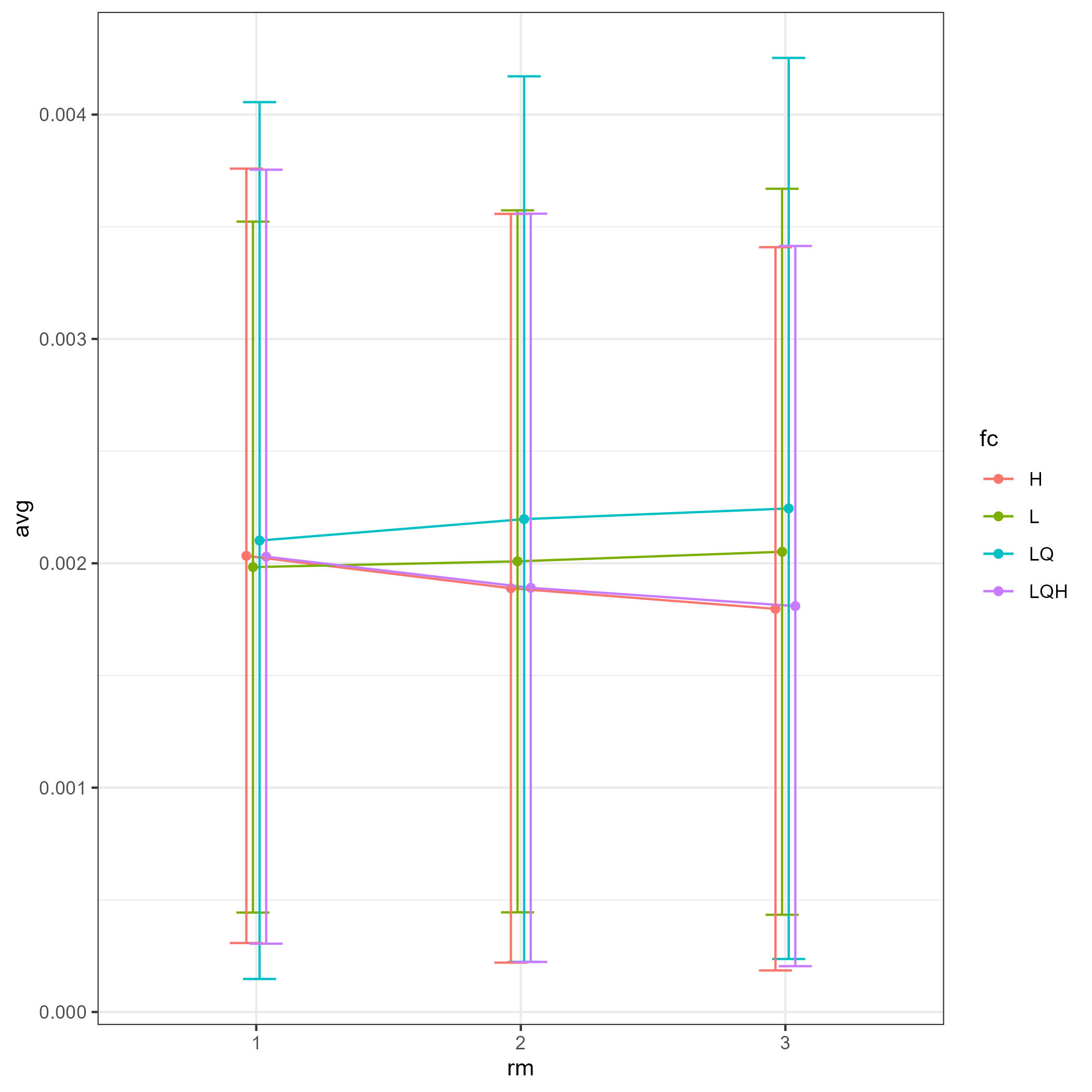

### auc_val.png

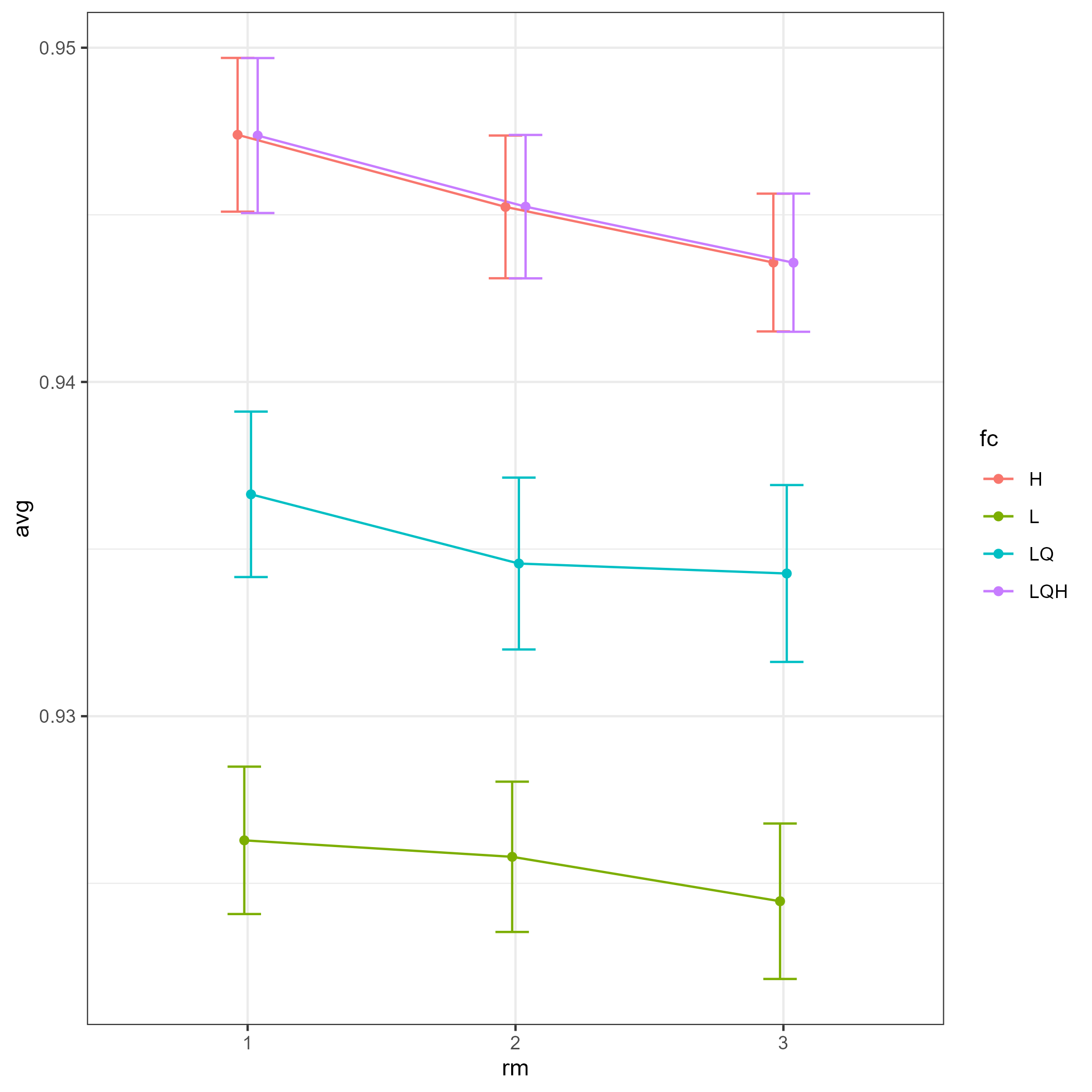

### bio01.png

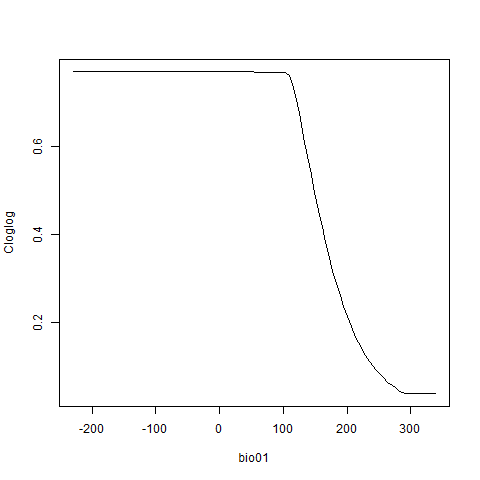

### bio02.png

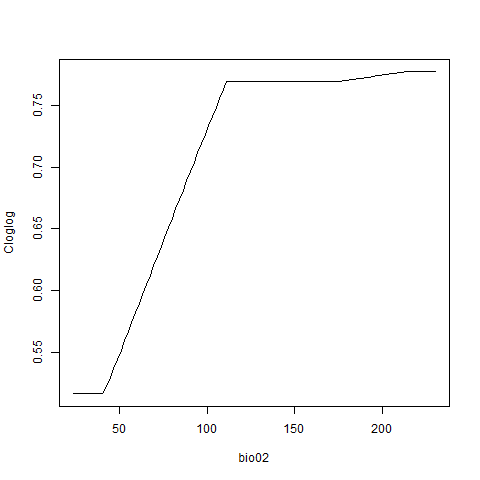

### bio03.png

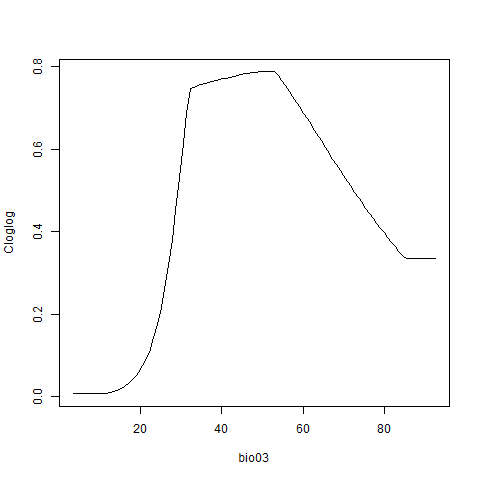

### bio04.png

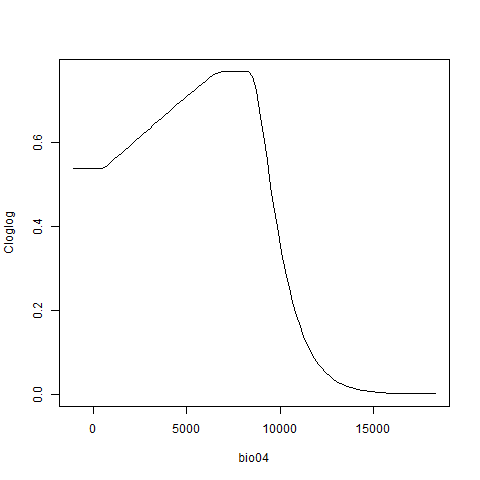

### bio05.png

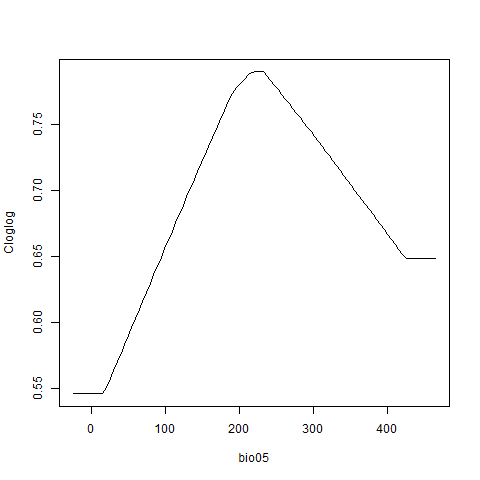

### bio06.png

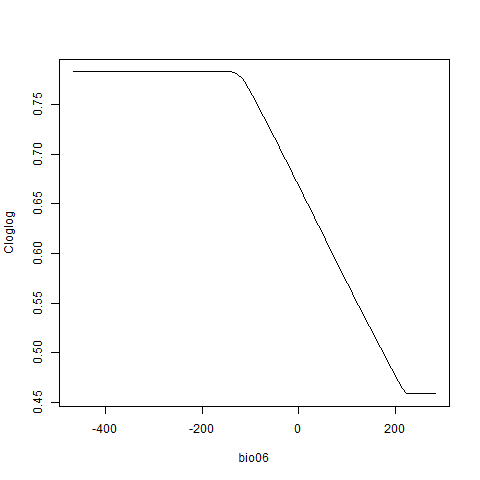

### bio08.png

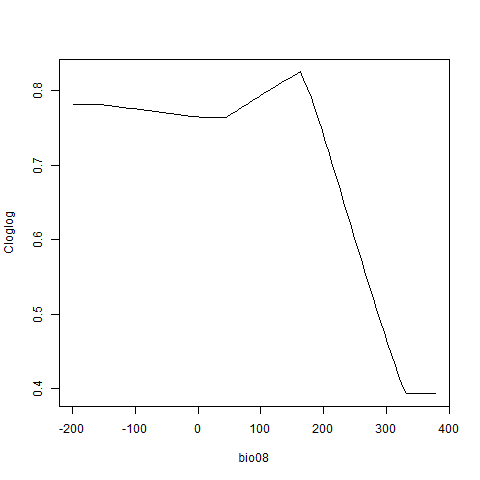

### bio09.png

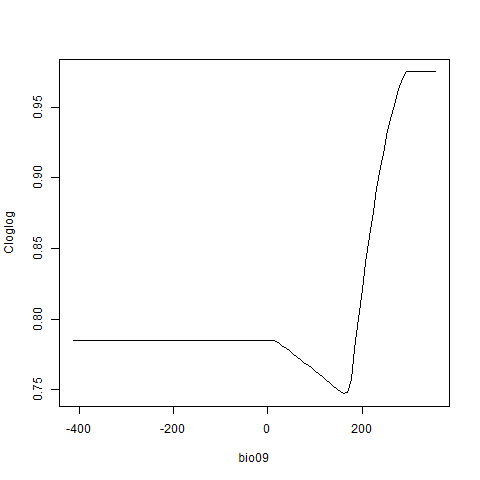

### bio12.png

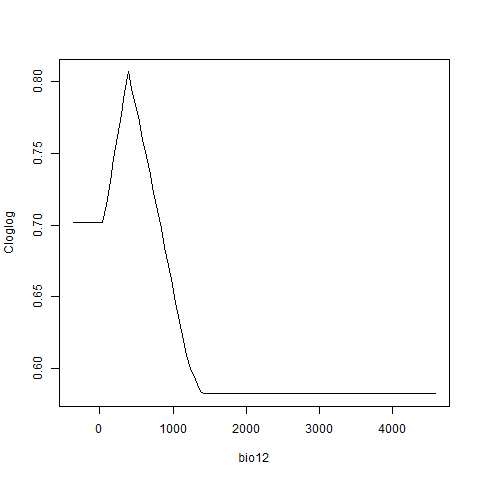

### bio14.png

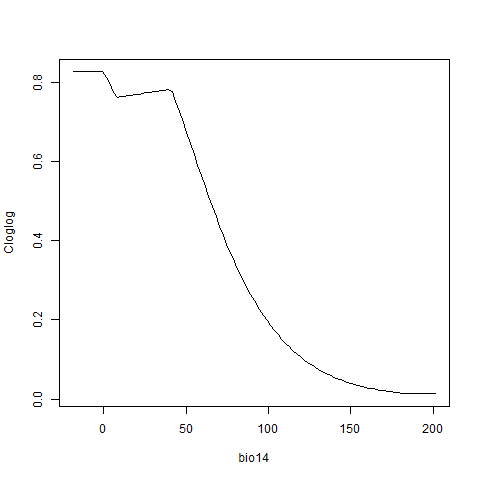

### bio15.png

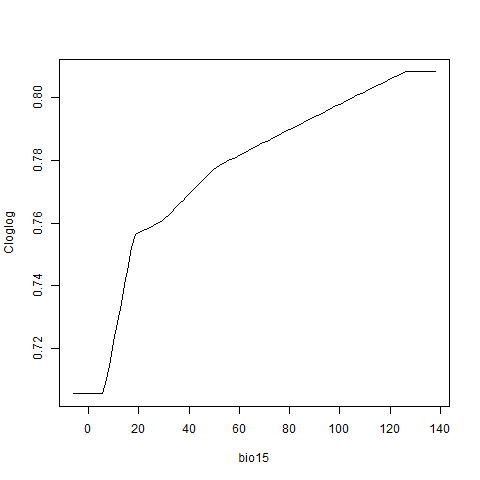

### bio17.png

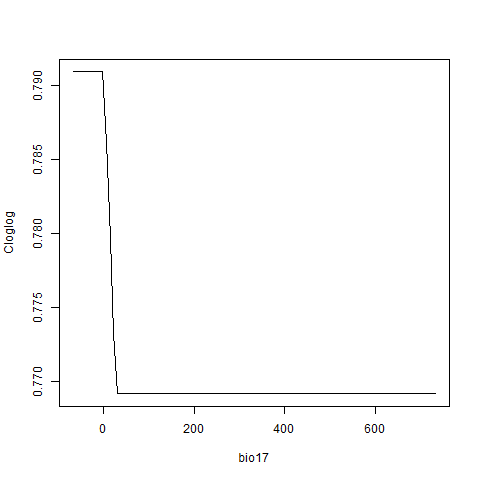

### bio18.png

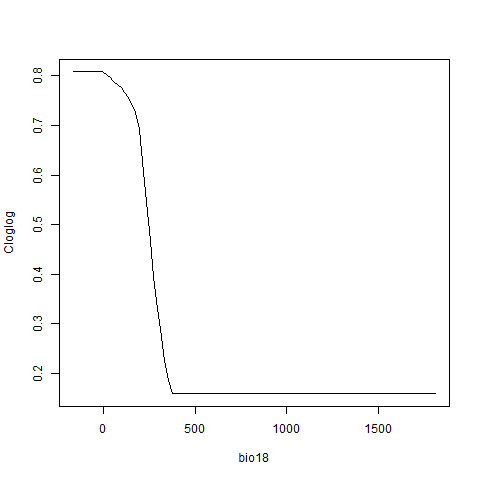

### bio19.png

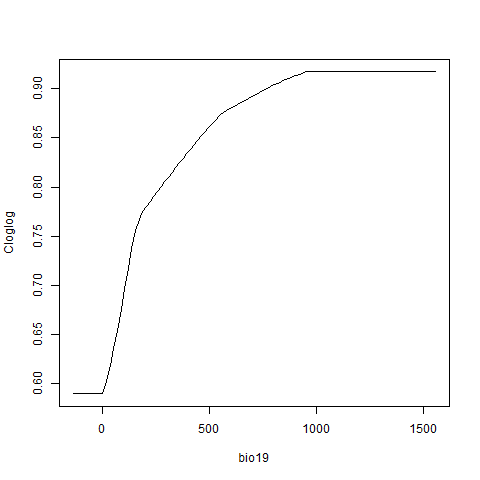

### delta_AICc.png

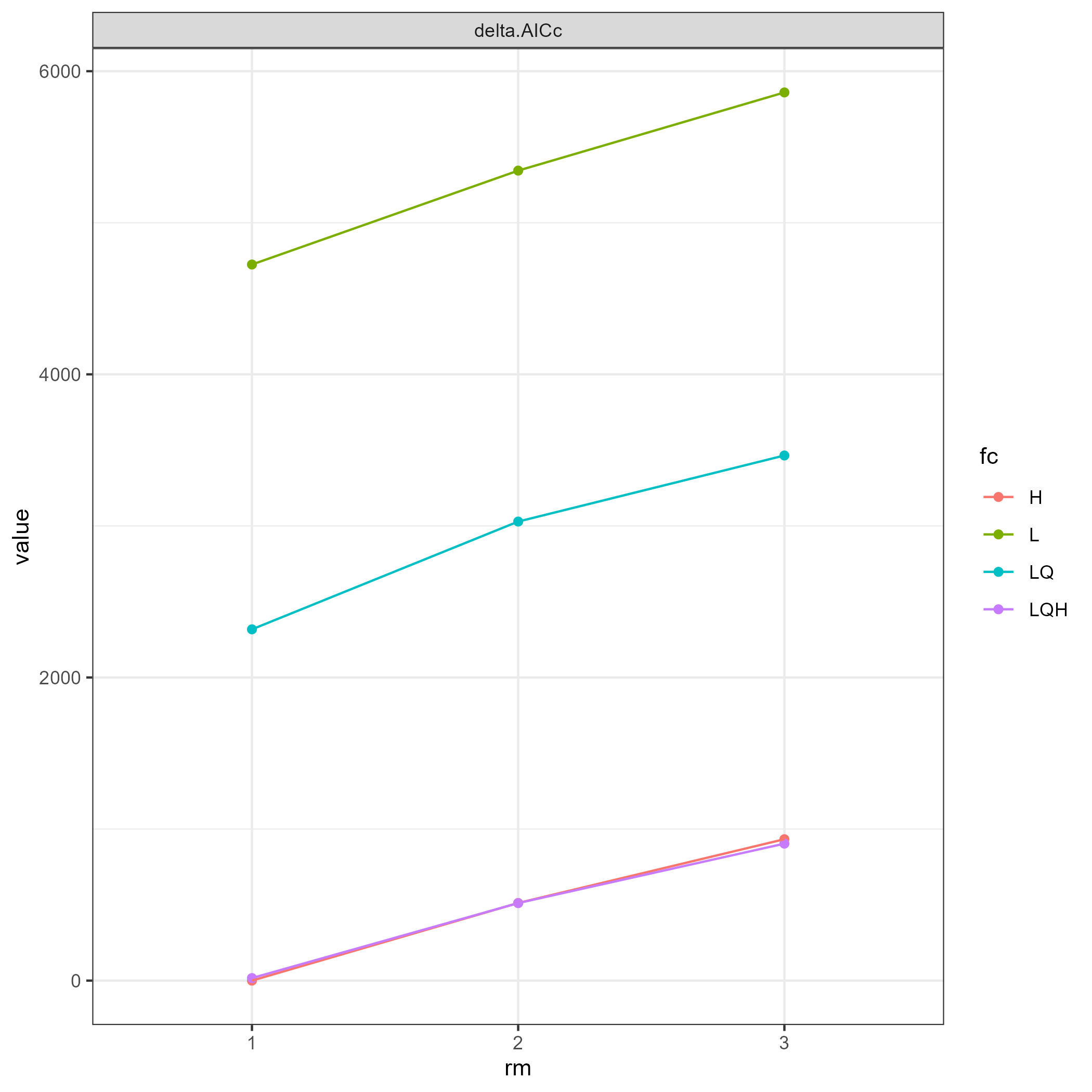

### or_10p.png

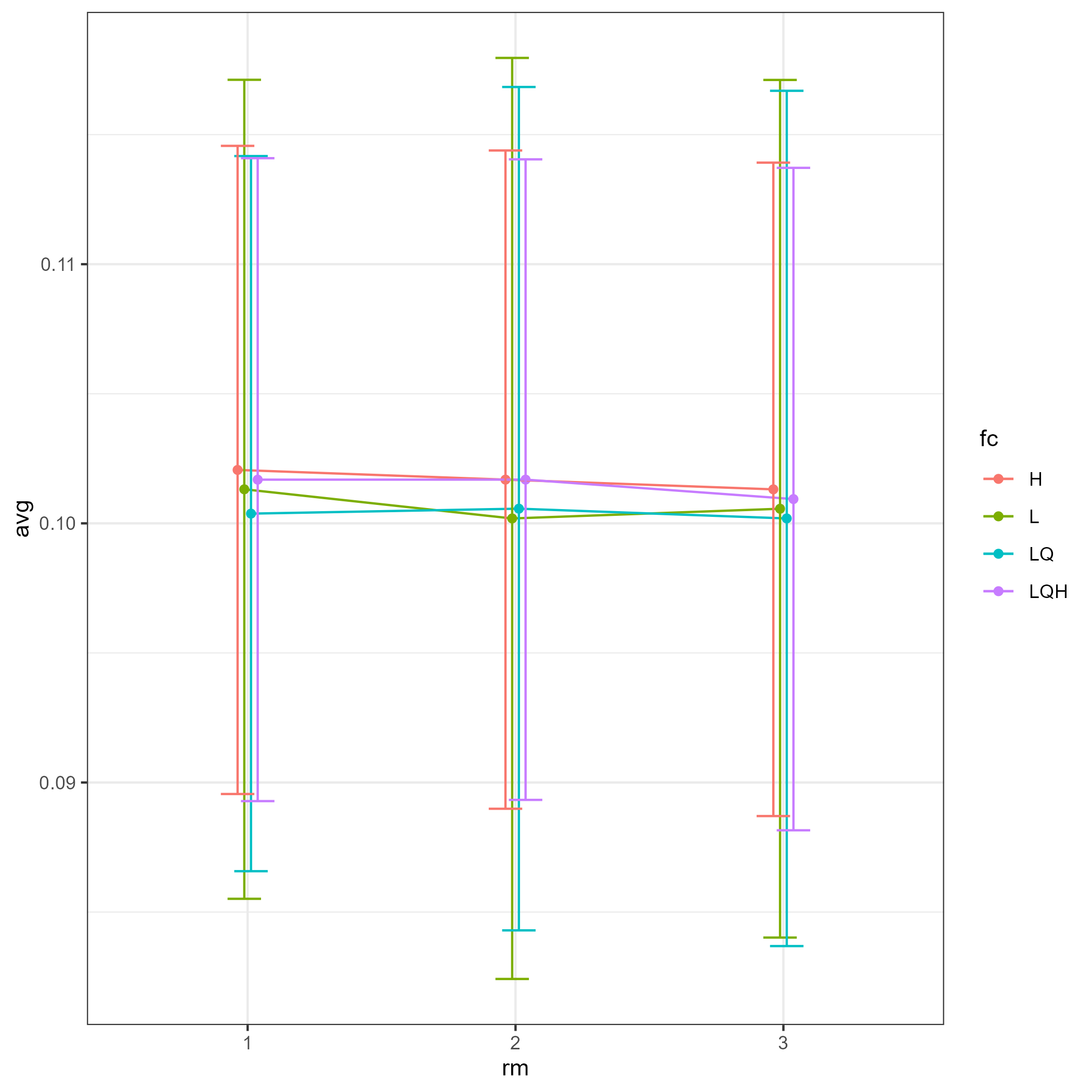

### or_mtp.png

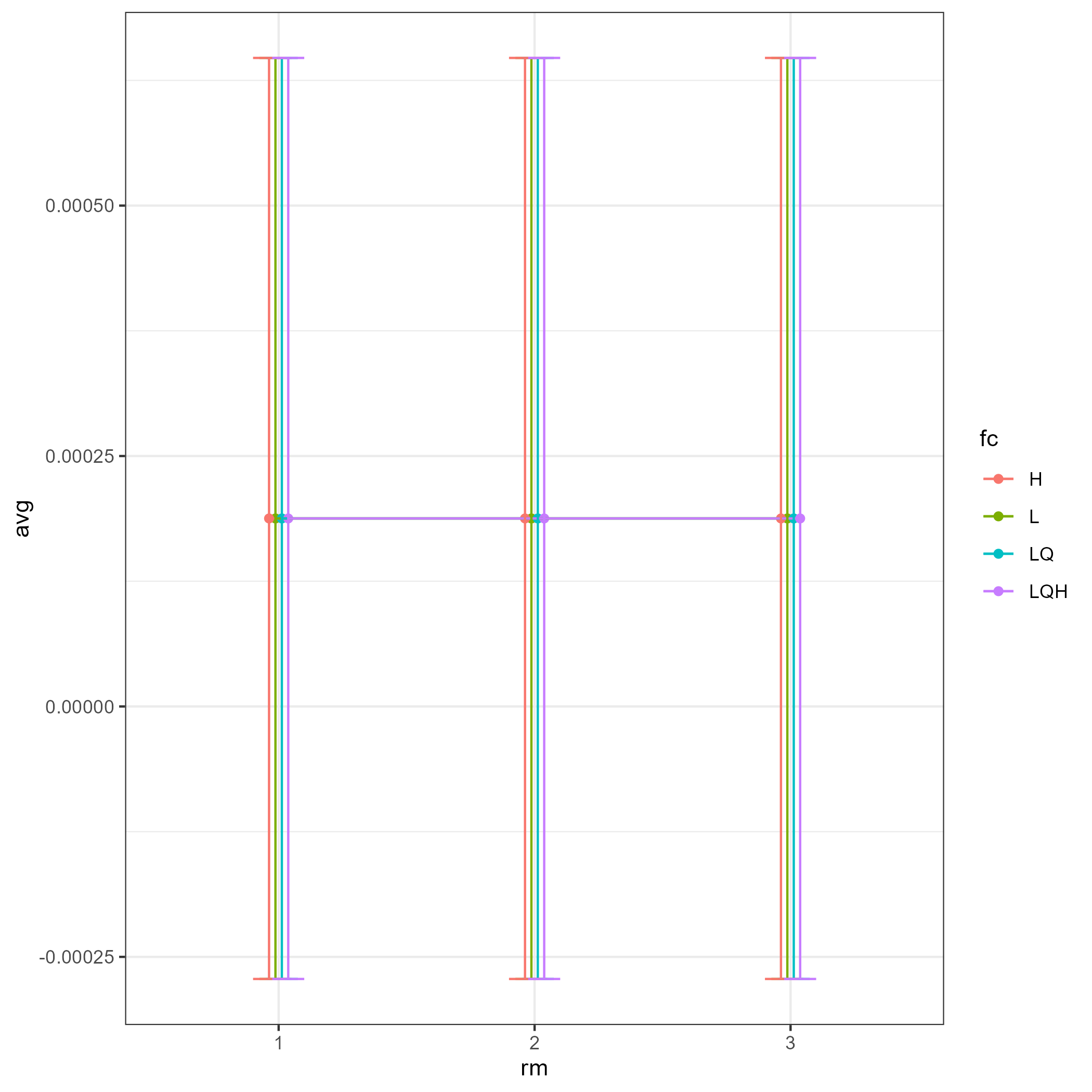
